# Goat Milk-derived Extracellular Vesicles modulate glial responses *in vitro* and attenuate neuroinflammation *in vivo*

**DOI:** 10.64898/2026.10.26.761897

**Authors:** Ana Santos-Coquillat, Paloma Lozano-Picazo, Eunate Iriarte-Urdiain, Jingjing Yang, Bernard Ucakar, Ariane Mwema, Marie Auquière, Anne des Rieux

## Abstract

Milk-derived extracellular vesicles (MEV) represent a naturally derived, potentially scalable platform for immunomodulation, yet their effects on central nervous system (CNS) glial inflammation at defined particle-to-cell ratios remain insufficiently characterized. Here, we investigated the immunomodulatory effects of goat milk-derived MEV in cellular and murine models of neuroinflammation. MEV-enriched preparations were obtained by sequential filtration, ultrafiltration, and size-exclusion chromatography, and characterized by scanning electron microscopy, nanoparticle tracking analysis, and dynamic light scattering. In LPS-stimulated BV2 microglia-like cells, MEV at 250 and 2500 particles per cell significantly reduced CD40 expression. In primary rat mixed glial cultures, microglia, astrocytes, and oligodendrocytes internalized MEV. At 250 particles per cell, MEV significantly reduced CD40 and IL1β gene expression at the mixed-culture level. In both cellular models, MEV also increased markers associated with an M2-like microglial phenotype under resting conditions. Following intrathecal administration in healthy mice, MEV did not induce significant changes in the inflammatory gene markers assessed at 8 h. In an LPS-induced mouse model of neuroinflammation, intrathecal MEV administration reduced CD40 protein expression towards control levels within 24 h and shifted the inflammatory marker profile towards a pro-resolutive state, as indicated by increased Arg1 levels and elevated CD206/CD86 and IL10/IL6 ratios. These findings identify goat milk-derived MEV-enriched preparations as potential modulators of CNS glial inflammation and support further investigation of their therapeutic potential.

## Introduction

Neuroinflammation is a component of numerous neurological diseases, including multiple sclerosis, Alzheimer’s disease, Parkinson’s disease, amyotrophic lateral sclerosis, traumatic brain injury, and spinal cord injury, where chronic immune activation drives the release of inflammatory molecules that damage neurons and glial cells and exacerbate disease progression [1, 2]. Current therapies for these conditions are largely limited to symptom management, and their efficacy is further constrained by the lack of drug delivery systems (DDS) capable of achieving optimized CNS accumulation [3]. Addressing this dual challenge, which requires identifying effective anti-inflammatory agents and developing delivery platforms suited to CNS administration, is essential to advance disease-modifying therapies and improve patient outcomes.

Among emerging candidates, extracellular vesicles (EV) have attracted particular interest. EV are nanometric, cell-derived particles that mediate intercellular communication by transferring proteins, lipids, miRNA, and DNA between cells [4]. Their nanoscale size, structural stability, and inherent biocompatibility make them an attractive natural nanocarrier platform, positioning EV not only as bioactive agents but also as a scaffold for engineered DDS. Milk is among the most extensively studied EV sources, with milk-derived EV (MEV) isolated from multiple species, including humans, pigs, cows, camels, and goats [5–9] and investigated as immunomodulatory nanomedicines of natural origin [10, 11]. MEV have demonstrated anti-inflammatory activity across several models, including suppressed cytokine expression in LPS-stimulated macrophages [12] and reduced serum MCP-1 and IL-6 in an arthritis model [13]. However, this activity appears highly dose-sensitive. Franzoni *et al.* reported that high MEV doses (60 μg protein equivalent) induced a pro-inflammatory M1-like phenotype in porcine monocyte-derived macrophages [14], and Hagey *et al.* showed that both particle concentration and EV subtype elicit distinct responses in recipient cells [15]. This variability underscores the need for dosing strategies that account for cell number rather than relying on total protein, protein concentration, or particle density alone. Here, we address this in our *in vitro* experiments by individualizing MEV quantity per recipient cell (henceforth, particle-to-cell dosing), enabling a more precise characterization of MEV’s dose-dependent immunomodulatory effects.

Considering all the possible sources, goat MEV stand up as an attractive candidate for immunomodulatory applications due to their wide availability, cost-effectiveness, and low allergenic potential [16] compared to, for example, cow milk. This choice also builds on a previously validated goat MEV isolation and characterization protocol established by the authors [17]. Despite this potential, no study to date has established an optimized, CNS-relevant particle-to-cell dosing strategy for goat MEV *in vitro* as a nanomedicine or DDS, a gap this study directly addresses. *In vivo*, goat MEV have shown therapeutic promise in diverse contexts, including neuroinflammation reduction via oral administration through the microbiota–gut–brain axis [18], as a nanovector for curcumin to reduce hepatic fibrosis in a mouse model [19], in pancreatic cancer therapy [20], or for tumor detection *in vivo* [21]. Most of these studies rely on oral [18, 22, 23] or intravenous administration [17, 19, 20], which limits their accumulation in the CNS, as it is known that most of the dose ends up in clearance organs [24]. In the present study, we evaluate direct delivery of MEV to the CNS via intrathecal (I.T.) injection into the cerebrospinal fluid.

Therefore, the objective of this work was to identify a goat MEV dose that maximizes anti-inflammatory activity while remaining within a safe, non-pro-inflammatory range, establishing a dosing framework relevant to their future development as a CNS-targeted drug delivery platform. We characterized MEV immunomodulatory properties in both non-activated and activated *in vitro* microglia and mixed glial cell models, evaluated their effects on the CNS of healthy mice following I.T. injection, and assessed their therapeutic efficacy in an LPS-induced neuroinflammatory model.

## Materials and methods

### Goat milk EV preparation

Goat MEV were prepared from fresh non-pasteurized milk (Chèvrerie De Volle Maan, Belgium) using previously developed methods by the authors [17, 25] (Figure 1A). Briefly, EV were obtained by differential centrifugation (3974 x*g,* 10 min, 4°C) and a microbial rennet combination for casein precipitation (Figure 1A). The resulting biofluid was centrifuged at 13000 x*g* for 30 min and 35000 x*g* for 15 min at 4°C with a high-speed centrifuge (Avanti® J-E, Beckman Coulter) to remove apoptotic bodies and microvesicles. The supernatant was then filtered through a 0.22 µm PES vacuum filter (Nalgene® 566-0020, Sigma-Aldrich, St. Louis, MO, USA) to remove larger components, and 70 mL were transferred to a Centricon® Plus-70 filter unit (regenerated cellulose, MWCO 30 kDa, UFC703008; Merck Millipore, Darmstadt, Germany), to concentrate the sample and remove components smaller than 30 kDa, followed by centrifugation at 3500 x*g* for 40 min at 4 °C. The filter unit was washed once with Dulbecco’s phosphate-buffered saline (DPBS, #14190144, Thermo Fisher Scientific, MA, USA) in the middle of the process. When the concentrate reached 1 mL, it was transferred to a size-exclusion column (SEC; Izon column qEV1, 35 nm, Lyon, France) to separate by nanometric size range, and fractions 7-11 were collected in a total volume of 2.8 mL using PBS as the elution buffer. EV protein content was quantified using Pierce™ microBCA Protein Assay Kit (Thermo Fisher Scientific, Massachusetts, USA) and NanoDrop™ 2000/2000c Spectrophotometer (Thermo Fisher Scientific, Massachusetts, USA). Although the isolation protocol yields an EV-enriched preparation rather than a fully pure EV population, as recommended by MISEV2023 guidelines when EVs and non-vesicular extracellular particles cannot be fully distinguished [26], for simplicity, this fraction is referred to as MEV throughout the manuscript.

**Figure 1.**
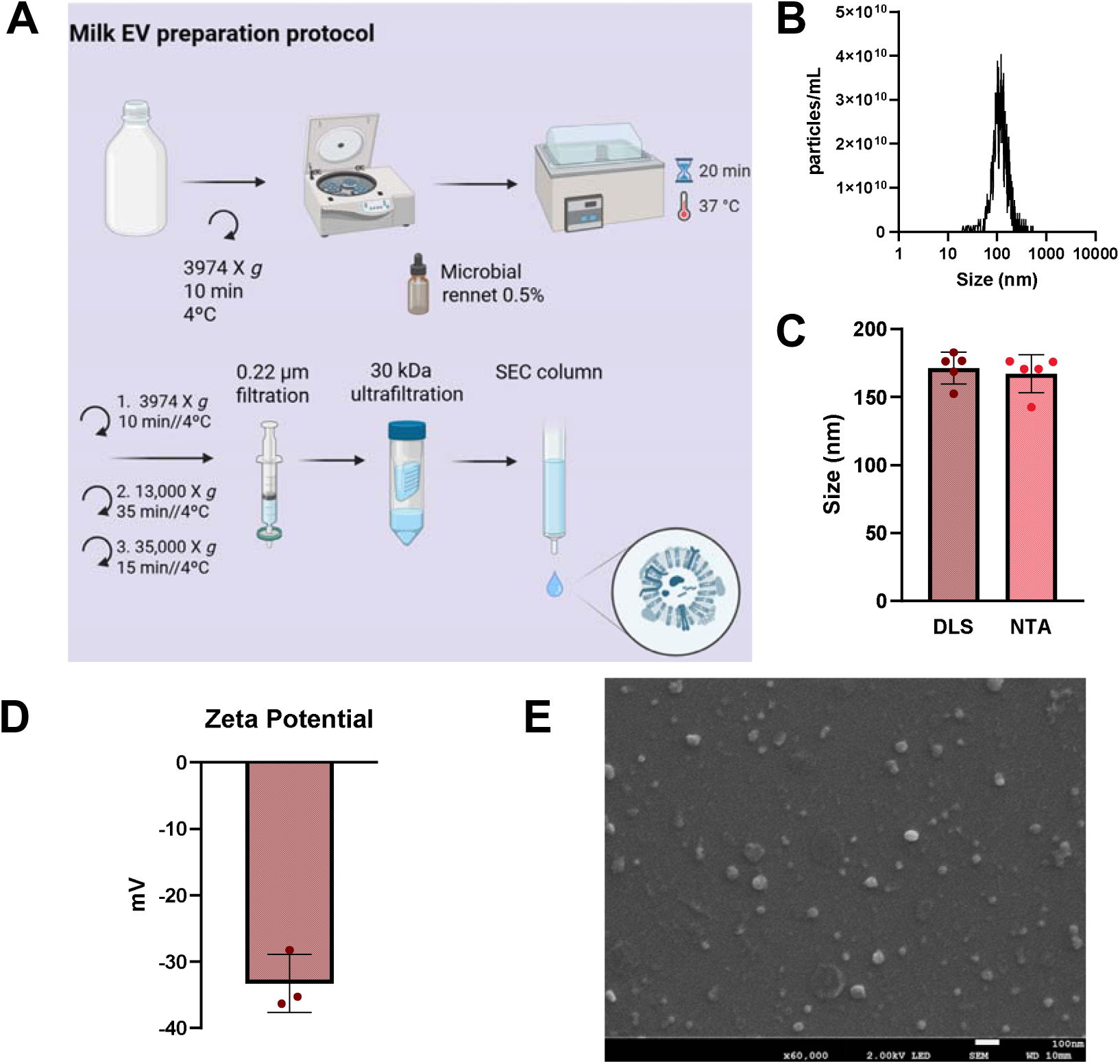
MEV preparation and characterization. A. MEV preparation workflow: differential centrifugation, microbial rennet incubation, subsequent centrifugations, 0.22 µm filtration, ultrafiltration (cutoff 30 kDa), and SEC purification. B. Particle size distribution and concentration by Nanoparticle tracking analysis (NTA), C. Size (nm) measurement by DLS and NTA (N=5), D. Zeta Potential (mV) determination (N=3). E. SEM images of MEV preparation at 60,000×; scale indicates 100 nm.

### Nanoparticle Tracking Analysis (NTA)

Real-time concentration (particles/mL) and size distribution of MEV were measured using NTA (ZetaView®, Particle Metrix, Inning am Ammersee, Germany). Five independent preparations were analysed. MEV were diluted in ultrapure water to a concentration ranging between 10^7^-10^8^ particles/mL. Sensitivity was set to 79 and camera shutter to 100. Measurements were averaged from particles counted in 11 positions for 2 repeated cycles.

### Dynamic light scattering (DLS) and Electrophoretic light scattering (ELS)

Size (N=5) and zeta potential (mV) (N=3) were measured using a NanoZS Zetasizer (Malvern Instruments Ltd., UK) and a DTS1070 disposable cuvette. Measurements were performed at 25°C, using protein as the material and water as the dispersant.

### Scanning Electron Microscopy (SEM)

Samples were diluted in DPBS at 1:10; a drop of the solution was placed on a Superfrost Plus Adhesion slide for 45 min, after which the excess liquid was removed, and 2.5% glutaraldehyde (Thermo Fisher Scientific, MA, USA) was added for 45 min, followed by 3x DPBS washing steps, dehydration (40%, 50%, 60%, 70%, 80%, 90%, and absolute ethanol) for 10 min each, and air-dried. The sample was coated with a 1 nm platinum layer by sputtering (Cressington sputter coater-208HR). SEM (JEOL JSM-7100F) was used with a 2 kV acceleration and working distance of 10 mm.

### Fluorescent labeling of MEV

MEV were labelled fluorescently with 50 µL of 16.9 mM Sulfo-Cyanine 5 NHS ester (SCy5) (Lumiprobe GmbH, Hannover, Germany) (90 min at RT at pH 8.5) starting from 500 µL of EV collected in fractions 7-11 (6× 10^11^ particles) [17]. Unbound dye was removed using an SEC column (Izon column qEV1, 35 nm).

### Cell culture

Murine microglial cells (BV2 cells) were cultured at 37 °C and 5% CO_2_ in high-glucose Dulbecco’s Modified Eagle Medium (DMEM, Thermo Fisher Scientific, MA, USA) with 10% Fetal Bovine Serum (FBS, Thermo Fisher Scientific, MA, USA), 100 U/mL penicillin, and 100 μg/mL streptomycin (#15140122, Sigma-Aldrich). StemPro™ Accutase™ (#A1110501, Thermo Fisher Scientific, MA, USA) was used for continuous passaging.

Cortical mixed glial cell (MGC) cultures were generated from Sprague-Dawley rat pups (postnatal day 0–2) as described previously [27, 28]. In brief, the brains were harvested, and the meninges, olfactory bulbs, and cerebellum were discarded. For tissue dissociation, the remaining brain material underwent enzymatic digestion with 40 μg/mL papain, followed by cautious mechanical trituration using G18 and G23 needles. The resulting cell suspension was prepared in DMEM (containing 4.5 g/L glucose, L-glutamine, pyruvate, 10% (v/v) FBS, and 100 U/mL penicillin and 100 μg/mL streptomycin) and passed through a 40 μm strainer. MGC were seeded onto poly-D-Lysine-coated 24-well plates (1 µg/mL) at a density of 1 × 10^6^ cells per well and maintained for 8 days, changing the media every two or three days.

### *In vitro* assessment of MEV effect on neuroinflammation

BV2 cells were seeded overnight into 24-well plates (2.5 × 10^5^ cells per well) and incubated with fresh culture medium containing 10 ng/mL LPS (From *E. coli*, serotype O55:B5, Sigma-Aldrich) and/or MEV at 250 or 2500 particles per cell (quantified by NTA), before being incubated for 8 h. MGC, cells were counted after 8 days of culture to determine cell number, and the medium was changed to fresh culture medium containing 10 ng/mL LPS and/or MEV at 250 or 2500 particles per cell for 8 h. Alternatively, MGC were incubated with 10 ng/mL LPS for 1 h, and then fresh medium containing MEV at 250 particles per cell was added and incubated for 24 h. Finally, 200 µL of TRIzol per well was added, and the cells were kept at -80°C for later assessment. Total RNA was extracted using TRIzol™ reagent (#15596026, Thermo Fisher Scientific) according to the manufacturer’s instructions [25]. Reverse transcription was performed using the GoScript^TM^ Transcription System (Promega, Madison, USA) from 800 ng of total RNA. qPCR was performed with GoTaq® qPCR Master Mix (Promega) and a STEP one PLUS instrument and software (Applied Biosystems, Foster City, USA). Data were analyzed with the ΔΔCt method using the 60S ribosomal protein L19 (RPL19) for mice or 60S ribosomal protein L13 (RPL13) for rat as reference genes. Primer sequences are given in Supplementary Table 1. For data normalization of MGC experiments, LPS expression levels from each independent experiment were set to 100 to cluster the data.

### Metabolic activity

BV2 cells were seeded overnight into 96-well plates (1.5 × 10^4^ cells per well) prior to being incubated with fresh culture medium containing 10 ng/mL LPS and/or MEV at 250 or 2500 particles per cell for 1 h or 4 h. PrestoBlue™ (Thermo Fisher Scientific, MA, USA) was used as per supplier’s instruction and incubated for 15 min before measurement using a SpectraMax ID5 (Molecular Devices) (Ex/Em 560/590 nm).

### In vitro uptake

BV2 cells or MGC were plated on 24-well plates (Corning® Costar, NY, USA) over glass coverslips at a cell density of 7 × 10^4^ cells or 6 × 10^5^ cells per well, respectively. 250 or 2500 MEV-SCy5 were incubated with BV2 or 250 MEV-SCy5 for MGC during 4 h.

BV2 cells were fixed with 4% PFA for 10 min, permeabilized with 0.1% Triton X-100 for 20 min, and incubated with Hoechst 33342 (Life Technologies; 1:1000) and Rhodamine-Phalloidin (ab235138, 1:1000 dilution, Abcam, Cambridge, UK) for 20 min at room temperature. Mean fluorescence intensity (MFI) was determined using ImageJ (Fiji) image analysis software by selecting regions of interest for each cell and measuring the mean gray value per channel.

MGC were fixed with 4% PFA for 15 min, permeabilized with 0.05% Triton X-100 in PBS for 20 min, and incubated with block solution (10% (v/v) FBS in PBS) for 1 h. Cells were then incubated at 4°C overnight with anti-GFAP (1:500; ab7260, Abcam), anti-Iba1 (1:500; 019-19741, Wako), or anti-NG2 (1:500; AB5320, Merck Millipore). Goat anti-Rabbit IgG (H+L) Cross-Adsorbed Alexa 488 (1:500, A1100, Invitrogen) was used as a secondary antibody. Nuclei were counterstained with Hoechst 33342 (Life Technologies). Dako Fluorescence Mounting Medium (Sigma-Aldrich, St. Louis, MO) was used to prepare the coverslips. Cells were observed under a confocal microscope (Zeiss LSM 980).

### Animal studies

All experiments were performed following the Belgian national regulation guidelines and EU Directive 2010/63/EU, following ARRIVE guidelines, and were approved by the ethical committee for animal care of the Faculty of Medicine of the Université catholique de Louvain (2024/UCL/MD/71). Seven-week-old C57BL/6JRj mice were obtained from Janvier Laboratories, France. Mice were housed in filter-top cages under artificial light (12-h dark/12-h light) and under controlled temperature (20–22 °C), with free access to food and drinking water.

### Intrathecal injection in healthy mice and LPS model

C57BL/6JRj mice (n=5 per group) were anesthetized using a rodent anesthesia system with vaporized isoflurane (2%) and 2*10^9^ particles (n=5) or PBS (n=5) were injected (20 µL/min) with a final volume of 10 μL using a 25-μL syringe and Remote Infuse/Withdraw Pump 11 Elite Nanomite Programmable Syringe Pump (Massachusetts, USA) in the intrathecal space between the groove of L5 and L6 vertebrae (tail flick), as described [29]. After 8 h, the brain was removed, processed, and analyzed by RT-qPCR. Total RNA was extracted from cerebellar brain tissue using TRIzol and processed using a Precellys Evolution Touch tissue homogenizer (Bertin Technologies, Montigny-le-Bretonneux, France). CD40, IL1β, CD206, CD86, Arg1, IL10, and IL6 were analyzed by RT-qPCR (Supplementary Table 1) as per the previous section.

For the LPS model, C57BL/6JRj mice (n=7-8 per group) were treated by intrathecal (I.T.) injection of ⁓2 *10^9^ particles or PBS, 1 h after intraperitoneal (I.P.) injection of LPS (0.5 mg/Kg) or saline. After 24 h, the brains were processed and analyzed by RT-qPCR, as previously explained, and ELISA.

For ELISA, midbrain tissues were homogenized in ice-cold lysis buffer containing 500 mM NaCl, 2 mM EDTA, 1 % Triton X-100, 0.5 % sodium deoxycholic acid, 0.1 % SDS, 50 mM Tris-HCl, and protease inhibitors. Samples were then sonicated using a Branson 450 digital sonifier® (Marshall Scientific, US) at 30 % amplitude for 10 s and subsequently centrifuged at 20,000 *g* for 10 min. Finally, the supernatants were collected for ELISA quantification. All measurements were normalized to protein content, determined using a Pierce™ BCA Protein Assay Kit (Sigma–Aldrich, USA). CD40 and IL-1β levels were quantified using Mouse CD40 ELISA Kit and Mouse IL-1β ELISA Kit (Invitrogen, Thermo Fisher Scientific, MA, USA) according to the manufacturer’s instructions.

### Statistical analysis

Statistical analysis was performed using PRISM 10.5.0 (GraphPad Software, CA, USA). Normality of data distribution was assessed for all datasets using the Shapiro-Wilk test. One-way ANOVA followed by Tukey’s multiple comparisons test was performed for normally distributed data (p > 0. 05), and the Kruskal-Wallis test followed by Dunn’s multiple comparisons test was applied to non-normally distributed data (p < 0.05). For the healthy mice *in vivo* experiments or dose comparison with MGC at 8 h, a t-test or Mann-Whitney test was performed after a normality test evaluation (Shapiro-Wilk). The number of experiments (N) and replicates (n) are indicated in the figure legends.

### Artificial intelligence tools

During the preparation of this manuscript, the authors used Claude Sonnet 5 (Anthropic) to improve the clarity and language of the text. All output was reviewed, edited, and verified by the authors, who take full responsibility for the content of this publication.

## Results and discussion

### MEV preparation and characterization

MEV had a median size of 170 ± 12 nm, regardless of whether we measured size by DLS or NTA (Figure 1B-C), a zeta potential of -33 ± 4 mV (Figure 1D), and 6 to 7 x 10^8^ particles per 1 µg of protein. SEM showed a size between 30 and 120 nm (Figure 1E). Using this protocol, we obtained 2.8 mL of product from 70 mL of purified volume, with a final yield of approximately 10^12^ total particles. This corresponds to around 100 mL of initial milk, as we can consider we lost 25-30% of the volume before the filtration steps, then reaching a yield of ∼1.1 × 10^10^ particles/mL of initial milk. This value closely matches the particle concentration reported for raw bovine milk (1.01 × 10^10^ particles/mL) [30]. This yield compares favorably to typical cell-culture-derived EV preparations, which commonly range from 10^8^ to 10^11^ particles/mL depending on cell type and culture format [31], underscoring milk as a scalable, high-yield EV source relevant to translational applications.

### Impact of MEV on microglia phenotype *in vitro*

Our objective was to study the impact of MEV on BV2 cells, a well-established microglial model, and how this effect was affected by the MEV/BV2 cell ratio. MEV uptake, cytotoxicity, and anti- and pro-inflammatory effects were studied *in vitro*. Two ratios of MEV/cell (high: 2500 MEV/cell and low: 250 MEV/cell) were tested to determine the required dose for a biological effect. Previous studies have tested from 20 to 20,000 particles per cell and found significant differences between doses when transcriptional effects were analyzed; importantly, EV cell source-specific responses were reported at the low range of doses [15]. Therefore, we selected two doses in the low range, with one order of magnitude between them.

We first confirmed that BV2 cells were able to internalize MEV. With both doses, cells were able to internalize MEV after 4 h (Figure 2A). This is consistent with previous data showing that goat MEV incubated with RAW 264.7 murine macrophages were internalized within a few hours [17]. The high dose increased the MFI values, with some cells taking up more than others. On the other hand, with the lowest dose, there was a more dispersed cytoplasmic accumulation that was difficult to distinguish at this magnification. Nevertheless, when quantified, the uptake was uniform across cells.

**Figure 2.**
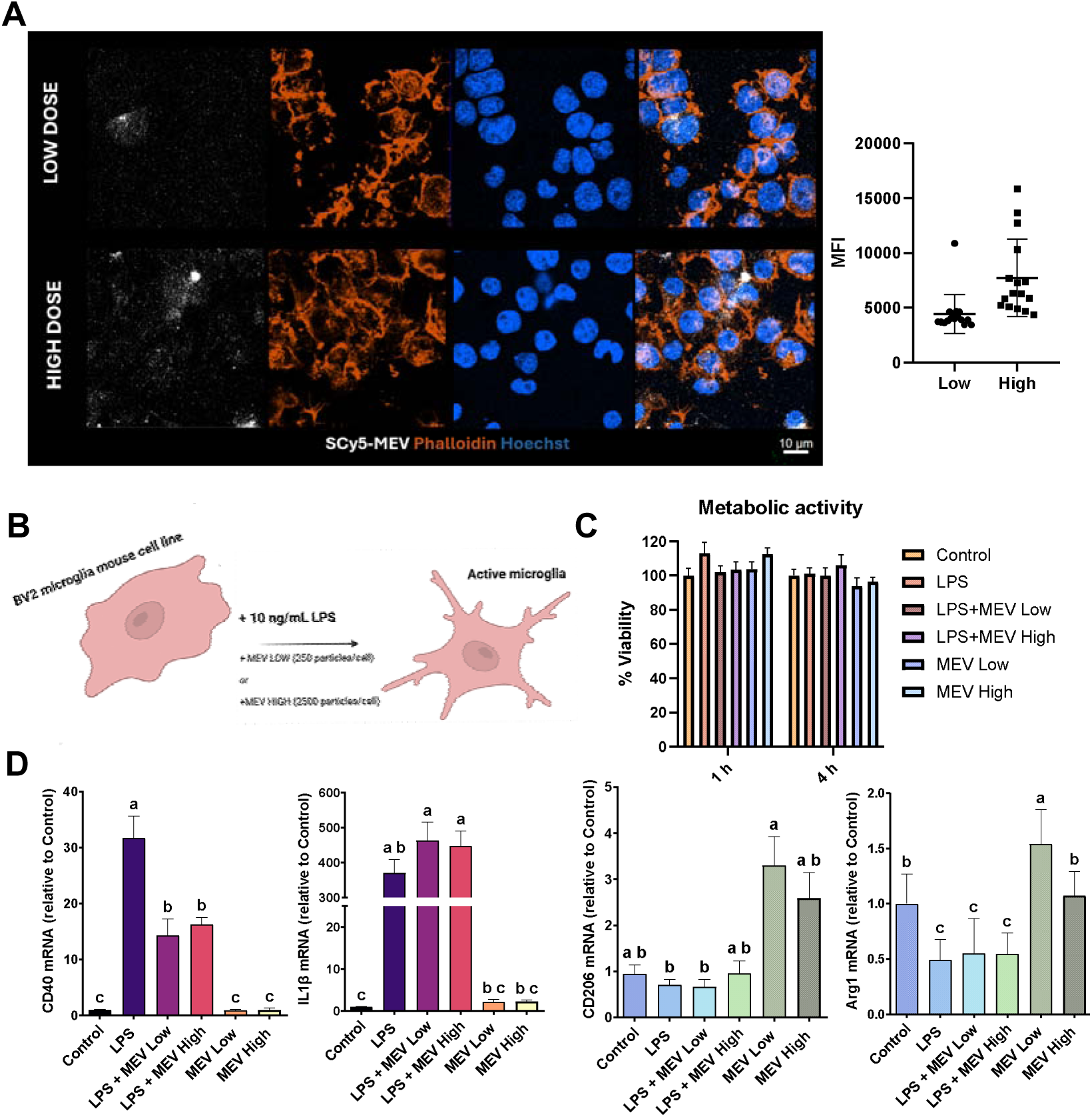
MEV uptake, cytotoxicity, and anti/pro inflammatory effects on mouse microglial-like cells. A. Confocal microscopy of BV2 uptake of SCy5-MEV at low (250 particles per cell) and high (2500 particles per cell) doses after 4 h of particle addition, SCy5-MEV in white, phalloidin in orange, and Hoechst in blue. Scale bar indicates 10 µm. Mean fluorescence intensity (MFI) values of SCy5-MEV uptake. B. Experimental setup for *in vitro* evaluation of MEV effects on neuroinflammation. Scheme prepared with BioRender ®. C. Metabolic activity of BV2 after 1 h and 4 h of exposure to MEV with or without LPS (10 ng/mL) (N=2, n=3-4). D. Determination of CD40, IL1β, CD206, and Arg1 by RT-qPCR after 8 h of exposure to LPS (10 ng/mL) and/or low or high dose of MEV (N=2, n=3-4); One-way ANOVA (CD40 and Arg1), Kruskal-Wallis (IL1β and CD206). Data with different superscript letters are significantly different. Error bars represent the Standard Error of the Mean (SEM).

Then, the influence of MEV and LPS exposure on BV2 cell metabolic activity was assessed after 1 h and 4 h (Figure 2B&C). No significant differences were observed, so neither LPS nor MEV, regardless of the dose, affected the metabolic activity of the cells. Franzoni *et al.* described that high MEV doses (600 µg of protein per 1×10^6^ cells, 0.6 µg protein/cell) can induce cell death in porcine macrophages after 24 h of exposure [14]. In our studies, we added ∼ 0.4 pg protein/cell or ∼ 4 pg protein/cell at the low and high doses, respectively. Moreover, 0.05, 0.5 and 5 µg of MEV protein/mL (corresponding to ∼ 0.67 pg/cell, ∼ 6.7 pg/cell, or ∼ 67 pg/cell, respectively, if we consider a volume of 200 µL in a 48 well plate) have been reported to be non-cytotoxic when incubated with RAW 264.7 macrophages for 24 h and 48 h [17] or up to 120 µg/mL of MEV (corresponding to 1200 pg/cell) on Human Skin Fibroblasts after 24 h [32]. This highlights the importance of evaluating cytotoxicity per cell type and raises the question of normalizing the EV dose/cell ratio to enable comparison across studies, an issue directly addressed by the particle-to-cell approach used here. Notably, even our high dose (∼ 4 pg protein/cell) remains several orders of magnitude below the cytotoxic threshold reported by Franzoni *et al.* (0.6 µg/cell) [14], and falls within the same order of magnitude as previously reported non-cytotoxic doses [17] once normalized per cell.

As human MEV has been reported to significantly decrease CD40 levels in LPS-activated BV2 cells [33] we studied the effects of goat MEV in the same model after 8 h of incubation. We observed that MEV effects were context dependent. Indeed, MEV significantly reduced CD40 expression without affecting IL1β, regardless of dose, in LPS-activated BV2 cells, while in resting cells, low-dose MEV increased Arg1 and CD206 gene expression without altering CD40 or IL1β. This suggests induction of an M2-like phenotype [34] only in non-activated cells (Figure 2D). At the high MEV dose, the inflammatory marker expression followed a similar pattern, indicating that increasing the dose did not enhance the anti-inflammatory effect of goat MEV. This confirmed that a higher MEV dose does not necessarily produce a greater biological effect. Overall, MEV promotes an M2-like phenotype in resting M0-like cells, but not in LPS-activated M1-like cells. Akinduro *et al.* also reported a significant reduction in IL1β after 24 h [33] which we did not observe at 8 h. Direct comparison is complicated by differences in dosing metrics (µg of protein versus particle-to-cell ratio, with cell number or total volume unspecified in [33]), in endpoints, as we studied it after 8 h while Akinduro *et al.* assessed it after 24 h, and in milk origin (goat vs. human). Biological source variability may further explain discrepancies across studies. Li *et al.* found that camel desert milk exosomes attenuated neuroinflammation and cognitive decline more effectively than non-desert milk exosomes [35], attributing this difference to a distinct proteomic profile enriched in AMPK pathway-associated proteins, suggesting that donor diet, lactation state, or species-specific milk composition are important variables to standardize in future MEV dosing studies.

To interrogate the effects of MEV in a more complex scenario where cellular communication between glial cells is maintained, rodent MGC were then treated with MEV. Microglial cells (Iba1+), astrocytes (GFAP+), and OPC (Oligodendrocyte progenitor cell; NG2+) were able to internalize MEV at the low dose (Figure 3A and Supplementary Figure 1A). Since MGC cultures have a higher cell density than BV2, more total MEV are present per imaging area even at the same particle-to-cell ratio, which may explain the higher apparent signal observed by confocal microscopy (Supplementary Figure 1A).

**Figure 3.**
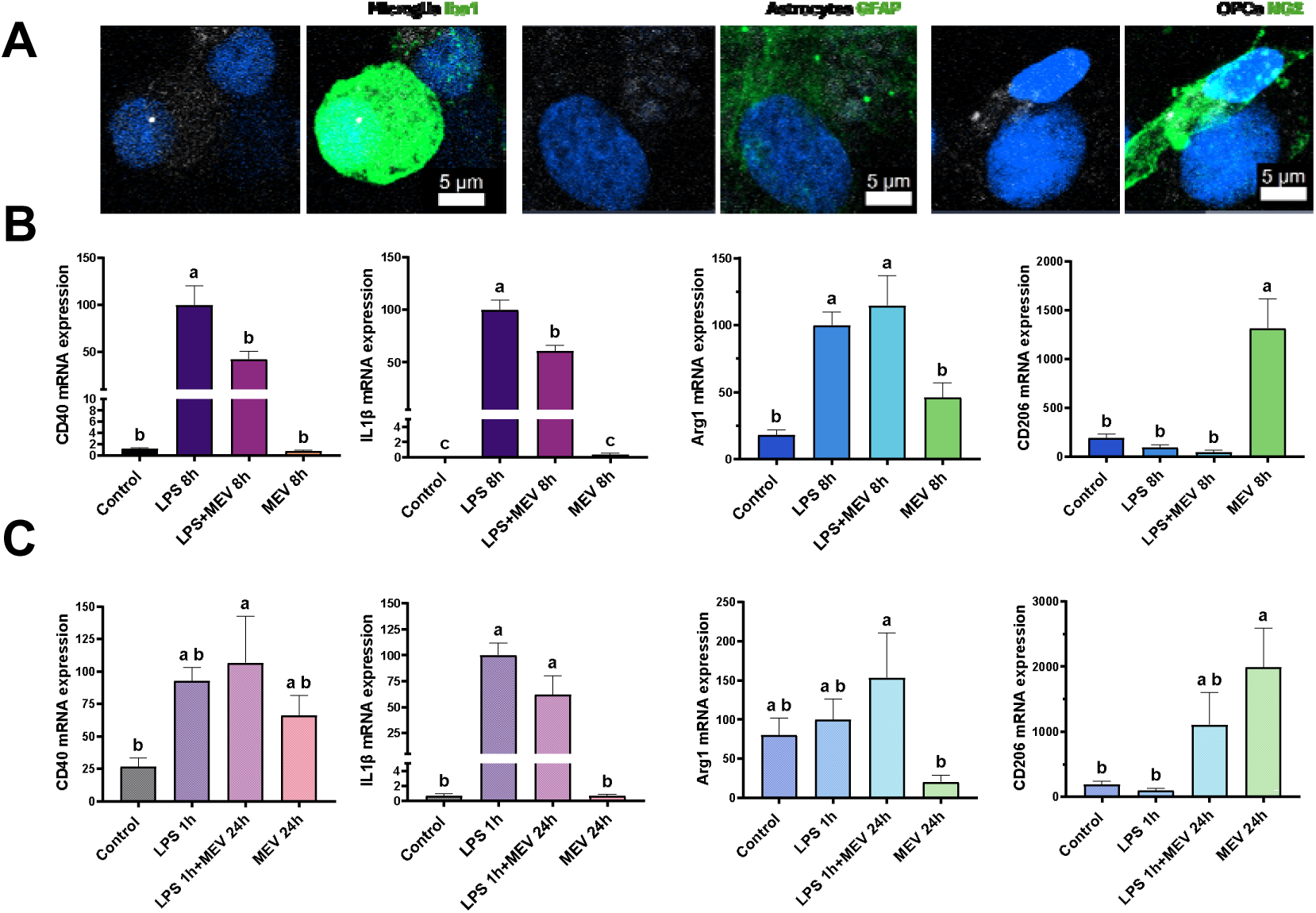
MEV uptake and anti/pro inflammatory effects on rat mixed glial cells (MGC). A. Magnification of the uptake by each type of cell (astrocytes, microglia or OPCs) with the low dose after 4 h. Scale bar indicates 5 µm. B. Determination of CD40, IL1β, Arg1, and CD206 by RT-qPCR after 8 h of exposure to LPS and/or low dose of MEV (N=3, n=3-4). C. Determination of CD40, IL1β, Arg1, and CD206 by RT-qPCR after 1 h of exposure to LPS and/or 24 h of MEV (N=2, n=3-8). CD206 after 24 h was analyzed by Kruskal-Wallis, and the rest by One-way ANOVA. Data with different superscript letters are significantly different. Error bars represent SEM.

In non-activated and activated MGC, gene expression of CD40 and IL1β was significantly reduced when MEV were added together with LPS on MGC after 8 h (Figure 3B). It has been described that after LPS exposure, there is an activation of microglial cells through TLR4, which secrete signaling molecules that induce astrocyte activation, which at the same time can regulate or activate the microglia population further [36]. Finally, crosstalk between both populations can induce toxicity to OPC cells [37]. Given that MEV was internalized by all three glial populations evaluated, including microglia, astrocytes, and OPCs, we hypothesize that MEV may attenuate neuroinflammation not only through a direct effect on microglia, but also by dampening the reciprocal microglia-astrocyte crosstalk that amplifies inflammatory signaling following TLR4 activation. This multi-cellular action could account for the more pronounced reduction in CD40 and IL1β observed in MGC, given that astrocytes also secrete IL1β [38], compared to the microglia-only BV2 model, where MEV reduced CD40 but not IL1β. Our result highlights the importance of *in vitro* evaluation in more complex cultures, such as MGC, as it can give a more realistic effect of nanomedicines before an *in vivo* evaluation, although the interpretation of results also becomes more challenging.

Related to Arg1 expression, Arg1 is significantly increased when LPS is added alone and with LPS plus MEV, and showed a non-significant two-fold increase in non-activated cells when compared to control. Arg1 upregulation upon LPS addition has already been described [39], and it does not occur when IFN-γ and TNF-α are used to polarize cells towards an M1-like phenotype.

Finally, CD206 was only significantly increased when the MEV were added to the non-activated MGC. As seen with BV2 cells (Figure 2D), MEV cannot induce an M2-like phenotype when added together with LPS for 8 h. A direct comparison of the low and high doses is presented in Supplementary Figure 1B, showing that the low dose had a higher anti-inflammatory profile when added to non-stimulated cells, while no significant differences were found between doses in activated cells, except for IL1β expression, which was higher at the high dose.

In view of these findings, we further explored whether the M2-like phenotype could be promoted with longer treatment times (24 h). MGC were therefore exposed to LPS for 1 h, the media was changed, and MEV at the low dose were added for 24 h (Figure 3C). CD206 was significantly increased when MEV were added alone; a similar, though non-significant, upward trend was also observed when MEV was combined with LPS, indicating that MEV could promote an M2-like phenotype regardless of LPS activation, although this effect was more robust in the absence of LPS. Arg1, on the other hand, showed a reduction in MEV-treated cells and an increase when LPS was added. We hypothesize that MEV alone may induce an early M2-like shift in Arg1 expression that is not sustained by 24 h.

LPS treatment increased IL1β levels, but when MEV were added, the expression was non-significantly reduced by up to 40% (p-value=0.17). MEV alone presented expression levels of IL1β similar to control. Also, CD40 increased with LPS alone with a non-significant upward trend (p-value=0.13) and significantly with LPS plus MEV (p-value=0.04). MEV alone were non-significantly increased more than two-fold compared to control (p-value=0.44). One explanation for not observing a reduction in CD40 after MEV addition is that protein-level regulation became more critical than gene expression by 24 h. Indeed, gene expression dropped 25-fold compared to 8 h, likely because the transmembrane protein was already present at the membrane. Qin *et al.* characterized CD40 protein expression in EOC13 microglial cells after 48 h of LPS exposure and found it to peak between 4-24 h [40]. Together, these results showed that MEV’s immunomodulatory effects on glial cells are dose- and time-dependent, with the clearest anti-inflammatory and pro-resolutive responses observed at low dose and in the absence of LPS. Several questions remain regarding the mechanisms behind these effects. Nevertheless, these mechanistic studies are beyond of scope of the study.

### Impact of MEV on neuroinflammation *in vivo*

To evaluate MEV’s safety and potential immunogenicity in healthy mice, MEV were injected *in vivo* directly into the CNS through an intrathecal (I.T.) injection. Other studies with EV from other sources have explored I.T. administration with promising results [41, 42]. This administration route is strictly constrained by a maximum injection volume of 10 µL in mice. Consequently, a dose of 9.5 × 10^10^ particles/kg (corresponding to 0.166 mg of protein/kg) was selected. Compared to established literature benchmarks for local administration, the particle count falls within the higher range, and this protein dosage sits at the lower end of the spectrum (average ∼ 0.5 mg/kg) [43]. These structural discrepancies and variations between protein weight and particle count directly reflect a broader, well-documented lack of methodological standardization *in vivo* EV dosing and clinical trials, where metric inconsistency remains a critical bottleneck [44].

Taking all of this together, we can consider that the dose selected was in the high range of the number of particles and the low range of protein quantity, indicating an effective MEV preparation protocol. I.T. injection of MEV in healthy mice showed that with a concentration of 2 * 10^9^ particles per mouse, there were no statistically significant changes in the gene expression of inflammatory markers (CD40, IL1-β, Arg1, CD206/CD86, IL10/IL6) in the brain when compared to the PBS-injected group after 8 h (Figure 4A). While a marginal, non-significant increase was visible for all genes, both pro-inflammatory and anti-inflammatory, the data demonstrated that this MEV dosage failed to induce active neuroinflammation in the CNS.

**Figure 4.**
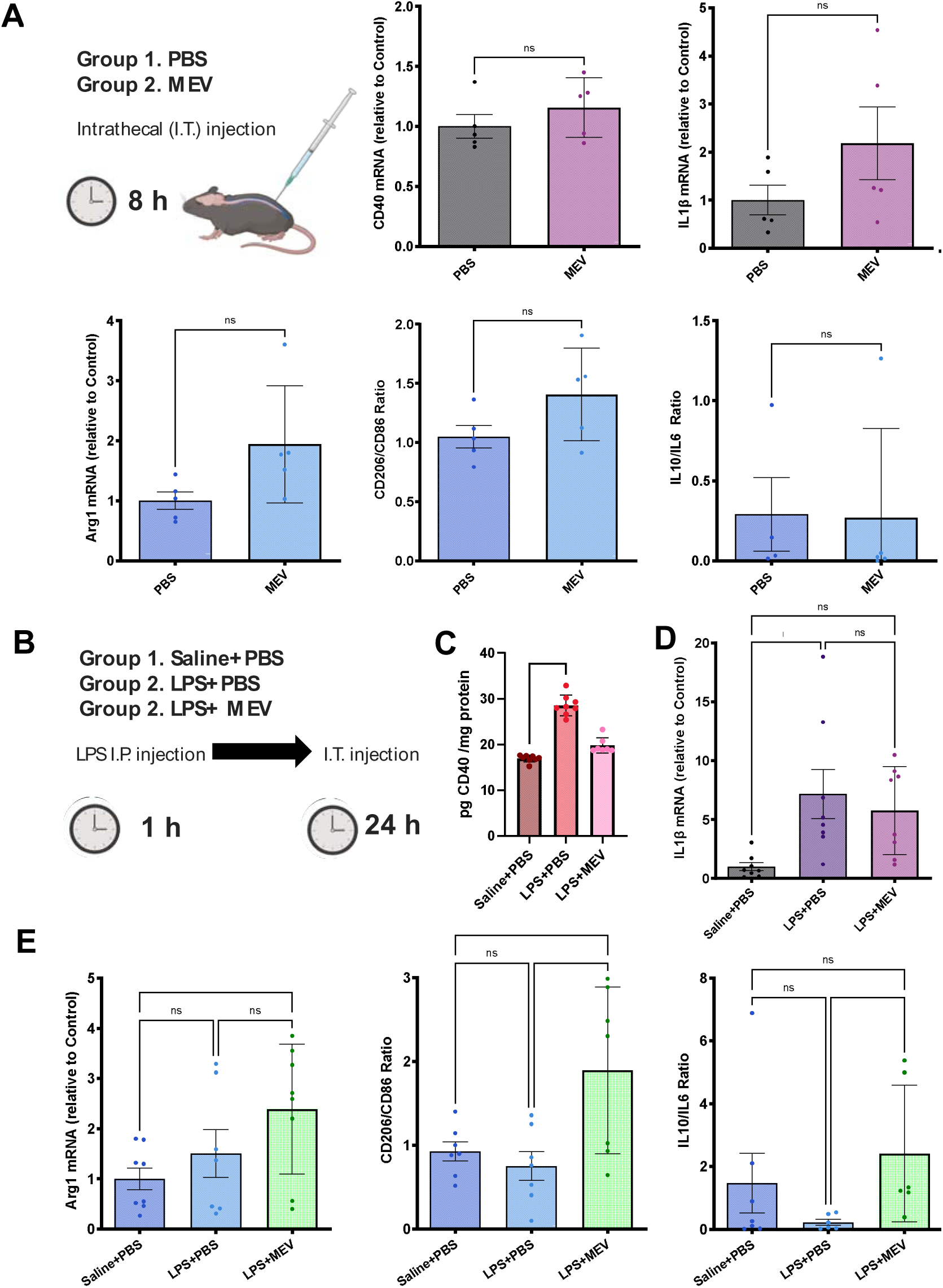
Evaluation of neuroinflammation modulation in healthy mice and a systemic inflammation model after MEV injection. A. Intrathecal (I.T.) injection in healthy mice of MEV or PBS (n=5). RT-qPCR determination of CD40, IL1-β, Arg1, CD206/CD86, and IL10/IL6 ratio expression in the brain 8 h after injection of the dose. T-test (IL1β, CD40, CD206/CD86) and Mann-Whitney (Arg1, IL10/IL6) B. LPS model of inflammation, n=7-8 per group; group 1: saline+LPS, group 2: LPS+ PBS, and group 3: LPS+MEV. C. ELISA determination of CD40 in the brain after 24 h. Kruskal-Wallis. D. RT-qPCR determination of IL1β expression in the brain after 24 h of injection of the dose. One-way ANOVA. E. RT-qPCR determination of Arg1, CD206/CD86, and IL10/IL6 ratio expression in the brain after 24 h of injection of the dose. One-way ANOVA (Arg1 and CD206/CD86) and Kruskal-Wallis (IL10/IL6). *p < 0.05, **p < 0.01, ***p < 0.001, ****p < 0.0001; ns = not significant. Error bars represent SEM.

We then wanted to evaluate the effects of this dose on modulating neuroinflammation; for that, we used an LPS model of systemic inflammation as a simple, cost-effective model to study neuroinflammation [45–47]. Here, LPS or saline was administered I.P. to the mice for 1 h, and then MEV or PBS was injected I.T., as in the healthy mice experiment (Figure 4B). As MEV incubation after LPS exposure in MGC for 24 h increased the expression of M2-like markers (Figure 3C), here the mice were sacrificed 24 h after LPS injection (23 h after MEV administration). Given the 24 h results with MGC (Figure 3C), where CD40 expression was reduced, we proceeded to quantify the CD40 protein levels in the brain, where we found them increased in the LPS+PBS group, whereas the LPS+MEV group showed CD40 levels comparable to the Saline+PBS control group (Figure 4C), indicating that MEV administration brought CD40 expression back toward baseline. IL1β mRNA showed a non-significant reduction of ⁓ 20% with MEV treatment compared to LPS alone (Figure 4D).

Overall, MEV treatment increased Arg1 levels significantly relative to healthy controls, the CD206/CD86 ratio increased significantly compared to both control and LPS groups and the IL10/IL6 ratio was also significantly increased compared to the LPS group (Figure 4E). Taken together, these results highlight the modulating effect of MEV in resolving inflammatory scenarios, which makes goat MEV an interesting nanomedicine for the CNS. Novel nanomedicines are still needed to finely tune neuroinflammation *in vivo* while avoiding an aberrant shift toward a pro-resolutive phenotype that promotes fibrosis or ECM deposition [34, 48].

Nevertheless, this study presents some limitations; the brain tissue was evaluated at a single time point (24 h) and therefore does not capture the long-term effects of MEV or the potential benefits of repeated or sustained dosing. Future *in vivo* work would also benefit from controlled-release strategies, for example, associating MEV with biomaterials [49], given the rapid clearance of EV *in vivo* and the importance of dose optimization observed.

As future perspectives, this study positions MEV as DDS candidates given that they can incorporate active drugs, such as RNA therapies, bioactive lipids, and hydrophobic compounds [10] and compared to MSC-derived EV, which are the most studied therapies [50] but require costly, labor-intensive cell culture and bioreactor scale-up. MEV can be obtained in large quantities directly from a readily available, low-cost biofluid, as demonstrated by the high particle yield achieved here from a small starting milk volume. Nonetheless, for these applications, the maintenance of their biological effects after drug association should be investigated.

## Conclusions

MEV modulate neuroinflammation by downregulating pro-inflammatory markers such as CD40 and/or IL1β *in vitro*, with effects observed in both microglial (BV2) and mixed glial cell (MGC) populations. In BV2 resting cells, MEV promoted an M2-like phenotype regardless of dose, while in MGC, this pro-resolutive effect was more consistent when MEV was administered alone at the low dose of 250 particles/cell, with a milder trend observed under LPS co-exposure. *In vivo*, intrathecal (I.T.) administration of MEV did not promote further inflammation in healthy mice, based on the inflammatory markers assessed after 8 h. In an LPS model of systemic inflammation, MEV treatment brought CD40 protein expression back towards control levels and promoted a pro-resolutive phenotype across multiple markers, including increased Arg1 and elevated CD206/CD86 and IL10/IL6 ratios. Ultimately, these findings support MEV as a naturally derived candidate for modulating CNS inflammatory responses.

## Acknowledgements

This work was supported by the F.R.S.-FNRS through the CDR J.0047.23. Ana Santos-Coquillat is grateful for financial support from F.R.S.-FNRS as *Chargée de recherches*. Eunate Iriarte-Urdain was supported by an Erasmus+ programme of the European Union. SEM images were obtained in collaboration with Ruben del Olmo from the VUB SUME group.

## Author Contributions

Ana Santos-Coquillat: Conceptualization, Methodology, Investigation, Writing – original draft, Writing – review & editing. Paloma Lozano-Picazo: Investigation, Methodology, Writing – review & editing. Eunate Iriarte-Urdiain: Investigation. Jingjing Yang: Investigation. Bernard Ucakar: Investigation. Ariane Mwema: Investigation. Marie Auquière: Methodology, Writing – review & editing. Anne des Rieux: Conceptualization, Methodology, Writing – review & editing, Funding acquisition.

## Data Availability Statement

The data that support the findings of this study are available on request from the corresponding authors.

