## Supplementary material for "Goat Milk-derived Extracellular Vesicles modulate glial responses *in vitro* and attenuate neuroinflammation *in vivo*": Suplementary material

**Supplementary Table 1.** Mouse and rat primer sequences.

| **Gene** | **Forward (5′-3′)** | **Reverse (5′-3′)** |
| --- | --- | --- |
| Mouse RPL19 | TGACCTGGATGAGAAGGATGAG | CTGTGATACATATGGCGGTCAATC |
| Mouse IL-1β | TCGCTCAGGGTCACAAGAAA | CATCAGAGGCAAGGAGGAAAAC |
| Mouse CD40 | GATTTGTGCCAGCCAGGAAG | AGAGAAACACCCCGAAAATGGT |
| Mouse CD206 | CCGTCTGTGCATTTCCATTC | TGCCAGTCAGTGGATCTTTGT |
| Mouse Arg1 | GGTTCTGGGAGGCCTATCTT | TGAAAGGAGCCCTGTCTTGT |
| Mouse CD86 | TTGTGTGTGTTCTGGAAACGGAG | AACTTAGAGGCTGTGTTGCTGGG |
| Mouse IL6 | ACAAGTCGGAGGCTTAATTACACAT | TTGCCATTGCACAACTCTTTTC |
| Mouse IL10 | GCCACATGCTCCTAGAGCTG | CAGCTGGTCCTTTGTTTGAAA |

| Rat RPL13 | GGCTGAAGCCTACCAGAAAG | CTTTGCCTTTTCCTTCCGTT |
| --- | --- | --- |
| Rat IL-1β | CCTGTGTGATGAAAGACGGCA | CTGCTTGAGAGGTGCTGATGT |
| Rat CD40 | TCATCCATGGACAAGCTGTGAA | CATTCAATGCTGGGGAAAGCAC |
| Rat CD206 | GAGGACTGCGTGGTGATGAA | CATGCCGTTTCCAGCCTTTC |
| Rat Arg1 | GAAGGTCTCTACATCACAGAAGAAA | CAAGGTCAACGCCACTGC |

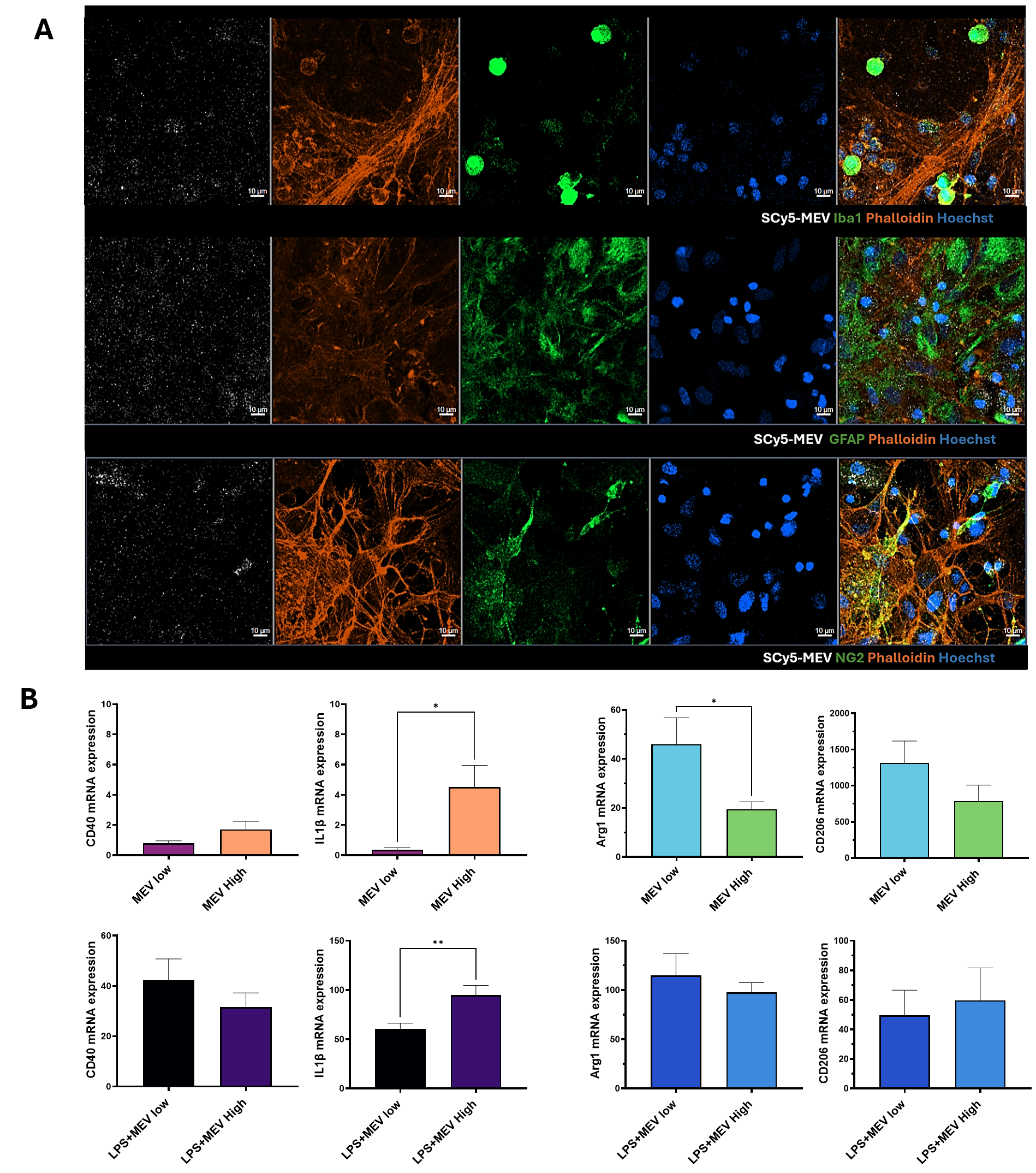

**Supplementary Figure 1. MEV uptake and comparison of anti/pro inflammatory effects of the low and high dose on MGC.** A. Confocal microscopy imaging of MGC uptake of SCy5-MEV at the low dose after 4 h, SCy5-MEV in white, phalloidin in orange, Hoechst in blue, Iba1, GFAP, and NG2 in green, respectively. Scale bar indicates 10 µm. B. Determination of CD40, IL1β, Arg1, and CD206 by RT-qPCR after 8 h of exposure to MEV low and high, without and with LPS (N=3, n=3-4). *p < 0.05, **p < 0.01. Error bars represent SEM.
